# Infected host competence overshadows heterogeneity in susceptibility in shaping experimental epizootics

**DOI:** 10.1101/2025.01.15.632224

**Authors:** Anna A. Pérez-Umphrey, Kate E. Langwig, James S. Adelman, Lauren M. Childs, Jesse Garrett-Larsen, Dana M. Hawley, Arietta E. Fleming-Davies

**Affiliations:** Department of Biological Sciences, Virginia Tech, Blacksburg, VA, 24061, USA; Department of Biological Sciences, University of Memphis, Memphis, TN, 38152, USA; Department of Mathematics, Virginia Tech, Blacksburg, VA, 24061, USA; Virginia Tech Center for the Mathematics of Biosystems, Virginia Tech, Blacksburg, VA 24061, USA; Department of Biology, University of San Diego, San Diego, CA, 92110, USA

**Keywords:** heterogeneity, host-pathogen interactions, prior exposure, susceptibility, transmission

## Abstract

The accelerated rate of disease emergence in recent decades underscores the need to understand conditions that promote or dampen epidemics. Theoretical models consistently show that epidemics are smaller in populations with higher among-individual heterogeneity in susceptibility. Experimental tests of these predictions are rare but critical for understanding how heterogeneity in susceptibility shapes epidemics in natural systems. We directly link data-parameterized models from previous dose response experiments in the house finch and *Mycoplasma gallisepticum* system to experimental epidemics in replicated aviary mesocosm flocks. We manipulated flock-level heterogeneity in susceptibility by seeding epidemics in flocks composed of either pathogen-naïve or previously exposed birds, which prior work showed have higher heterogeneity in susceptibility relative to pathogen-naïve populations. We tracked epidemics for over two months, combining empirical data and stochastic compartmental models to address how heterogeneity in susceptibility changes epidemic severity. Consistent with previous work, estimates of heterogeneity in susceptibility based on coefficients of variation were higher for flocks given prior pathogen exposure relative to pathogen-naïve flocks. However, in contrast with prior work on individually-housed birds which showed relatively homogeneous susceptibility for pathogen-naïve birds, the pathogen-naïve flocks in this study were better described by heterogeneous, rather than homogenous, models of susceptibility. This suggests that flock-level epidemics captured sources of heterogeneity absent in controlled experiments, such as transmission heterogeneity. Finally, although prior exposure conferred protection from disease at the individual level, we did not detect predicted effects of prior exposure and its associated flock-level heterogeneity on prevalence. However, our ability to detect effects of prior exposure on flock-level prevalence was obscured by unexpected variation in the competence of the initially pathogen-naïve index birds that seeded each epidemic. This variation in infectiousness among index birds significantly predicted flock-level prevalence, with low index bird infectiousness contributing to the absence of detectable epidemics in two of the three naïve flocks. Our stochastic simulations generated a wide range of prevalence outcomes for small epidemics over the timescales examined, further underscoring the challenges of measuring transmission dynamics in naturalistic settings, where unexpected variation in host traits such as competence can obscure other factors of interest.

**Open research statement:** Data are not yet provided. Data and code will be permanently and publicly archived in the Virginia Tech Data Repository if the paper is accepted for publication

## Introduction

The accelerated emergence and spread of pathogens in both human and wildlife populations (Jones et al. 2008; Rosenberg 2015; Smith et al. 2014) underscores the urgent need to better characterize the conditions that promote or dampen epidemics and their severity. A growing body of work suggests that key traits of hosts in a population are often highly heterogenous in nature, and this complexity can have considerable epidemiological influence (Lloyd et al. 2020; Gomes et al. 2022; Elie, Selinger, and Alizon 2022; Aalen et al. 2015). Some sources of host heterogeneity have been well-studied, such as transmission-relevant traits (VanderWaal and Ezenwa 2016; Vazquez-Prokopec et al. 2016), including heterogeneity in infectiousness (Woolhouse et al. 1997; Lloyd-Smith et al. 2005; White et al. 2020) and host competence (Gervasi et al. 2015; Cortez and Duffy 2021), or contact rates and exposure (Collins and Govinder 2014; Paull et al. 2012; Hamede et al. 2012). This body of work has consistently highlighted the importance of incorporating certain types of heterogeneity into disease models to accurately predict outbreaks and apply realistic control measures (Collins and Govinder 2014; Lloyd-Smith et al. 2005; Paull et al. 2012). While less well-studied, how heterogeneous a population is in its susceptibility to a given pathogen (defined here as the probability of infection given exposure) can also have far-reaching downstream epidemiological (Rodrigues et al. 2009; Rose et al. 2021; Hawley et al. 2024; Langwig et al. 2017; Dwyer, Elkinton, and Buonaccorsi 1997) and evolutionary (Fleming-Davies et al. 2015; Read et al. 2015) consequences. However, to date, there have been no opportunities to experimentally test how alterations in the degree of heterogeneity in susceptibility in a population alter epidemic outcomes.

Heterogeneity in susceptibility among individuals can stem from intrinsic factors (e.g., genetics, age class) but can also be induced or augmented by prior pathogen exposure, whether that be through vaccination or natural infection (Gomes et al. 2014). Prior pathogen exposure reduces the mean susceptibility of a population while also often increasing the variance by generating incomplete or variable immune protection across individuals. Even where exposure dose can be controlled (i.e., vaccines), the degree and duration of protection generated can vary substantially across individuals, inducing population-level heterogeneity (Le et al. 2021). While the degree of heterogeneity in susceptibility can be challenging to measure empirically because it is inherently a population-level metric, dose-response experiments that measure infection probability over a series of controlled challenge doses (Gomes et al. 2014; Haas, Rose, and Gerba 2014; Ben-Ami, Ebert, and Regoes 2010) allow direct quantification of population-level heterogeneity in susceptibility that are not confounded by other sources of host variation. Using these approaches, two studies to date demonstrated that vaccination against or prior exposure to a pathogen augmented heterogeneity in susceptibility: rainbow trout vaccinated for infectious hematopoietic necrosis virus (IHNV) show significantly higher population-level heterogeneity in susceptibility than unvaccinated trout (Langwig et al. 2017), and house finches (*Haemorhous mexicanus*) with prior exposure to either a low or high dose of their naturally-occurring pathogen *Mycoplasma gallisepticum* show higher heterogeneity in susceptibility relative to pathogen-naïve birds (Hawley et al. 2024).

Population-level heterogeneity generated by prior pathogen exposure is particularly important because theoretical models consistently show that heterogeneous susceptibility dampens the size of epidemics (Langwig et al. 2017; Ben-Ami, Ebert, and Regoes 2010; Katriel 2012; Gomes et al. 2014; Gomes et al. 2022; Dwyer, Elkinton, and Buonaccorsi 1997), in part because the most susceptible individuals become infected first (“cohort selection”) while the most resistant individuals remain (Katriel 2012; Gomes et al. 2014; Langwig et al. 2017; Hawley et al. 2024). For instance, in an SIR model of vaccinated versus unvaccinated rainbow trout, Langwig et al. (2017) found that when controlling for the expected mean reduction in susceptibility from vaccination, population-level heterogeneity in susceptibility alone still resulted in a 35% reduction in outbreak size relative to homogeneous – or non-vaccinated – populations. Further, using experimental data on previously exposed house finches to parameterize SIR models, epidemics were substantially reduced (> 55%) in populations with heterogeneous relative to homogenous susceptibility, even when controlling for differences in mean susceptibility (Hawley et al. 2024). Importantly however, how heterogeneity in susceptibility shapes epidemics in natural settings, where additional sources of heterogeneity are present, has not been empirically tested. Here we use the house finch – *Mycoplasma gallisepticum* system to directly link data-parameterized models to experimental epidemics in aviary mesocosm flocks.

The house finch is a common backyard North American songbird that experiences seasonal epidemics of a bacterial pathogen (*Mycoplasma gallisepticum*; MG), which causes severe conjunctivitis (Ley, Berkhoff, and McLaren 1996). Infection-induced mortality in this system is largely indirect, primarily due to increased predation of infected birds in the wild (Adelman, Mayer, and Hawley 2017), but birds in captivity (in the absence of predators) typically recover. Birds that recover from prior infection have incomplete acquired protection, with antibody levels further waning over time since initial exposure (Fleming-Davies et al. 2018) – characteristics that augment heterogeneity in susceptibility to reinfection (Hawley et al. 2024). But, importantly, and as in any natural disease system, multiple other sources of transmission heterogeneity exist, each with a differential degree of influence on epidemiological outcomes (VanderWaal and Ezenwa 2016). For instance, in this system, characteristics such as the degree and duration of infectiousness (as measured by pathogen loads) can vary substantially among infected house finches (Hurtado 2012; Leon, Fleming-Davies, and Hawley 2019). Contact rates or exposure risk can also be highly heterogenous. Because MG transmission is primarily linked to the use of bird feeders that act as fomites (Dhondt et al. 2007), foraging behaviors (e.g., time spent at bird feeders) are correlated with pathogen transmission and acquisition (Adelman et al. 2015), which covary with other factors such as social dominance and ambient temperature (Adelman et al. 2013; Hawley et al. 2007; Teemer and Hawley 2024). In this way, some birds may act as “super-spreaders” and others as “super-receivers” (Adelman et al. 2015).

The well-characterized MG-house finch system is poised to examine heterogeneity in susceptibility and its epidemiological influence under semi-natural conditions. Here, we experimentally test whether prior exposure-induced population heterogeneity in susceptibility dampens the size of experimental epidemics in free-flying aviaries, which naturally harbor other sources of transmission heterogeneity. Unlike past experimental dose-response studies, our use of a mesocosm experimental design examines heterogeneity in susceptibility in a context where it is, by definition, broader: because exposure dose and transmission cannot be controlled for under natural conditions, in this context heterogeneity in susceptibility is better defined as heterogeneity in the likelihood of hosts to become infected. We do this by initiating epidemics in mesocosm flocks comprised of house finches with experimentally manipulated exposure histories: either no prior exposure (pathogen-naïve) or prior exposure to a low dose of MG, which past work showed significantly augments population-level heterogeneity in susceptibility (Hawley et al. 2024). We longitudinally track epidemics over more than two months and combine empirical data and compartmental models to address how prior exposure-induced heterogeneity in susceptibility (i.e., host risk of infection) influences epidemic severity.

## Methods

### Experimental design

To test how prior exposure to MG alters epidemic dynamics, we set up experimental flocks comprised of birds that varied in their exposure history and seeded experimental epidemics of MG using three “index” birds per flock (Figure 1). All birds in the study were wild-caught and confirmed immunologically pathogen-naïve before the experiment (Appendix S1:Section S1). A total of 84 “flockmates” were randomly assigned to either the prior exposure (n = 42) treatment or no prior exposure (hereafter, referred to as “naïve”; n = 42), which received a sham inoculation. Index birds were treated identically and were all initially pathogen-naïve to control for potential effects of prior exposure on infectiousness (Leon et al., 2025). Sex ratios were made as even as possible within treatment groups and flocks, although some incidental mortality unrelated to infection (see sample sizes in Appendix S1:Table S1) and the presence of males with female-typical plumage before molting resulted in a skewed sex ratio for one flock (Table 1; note that this had no apparent statistical effect [Appendix S1:Section S2]). Birds were kept in individual cages during the controlled prior exposure treatments and then allowed to recover from this initial pathogen exposure for 38 days. Then, birds were placed into one of six flocks (three per treatment) for experimental epidemics, with 17 birds per flock (14 flockmates from the same treatment group [prior exposure or naïve] plus three index birds). Flocks were established in six large replicate aviary units (Appendix S1:Section S1), which are partially exposed to the environment so that animals experience ambient conditions. The experimental epidemics began when index birds were inoculated and re-released back into their flocks.

**Figure 1.**
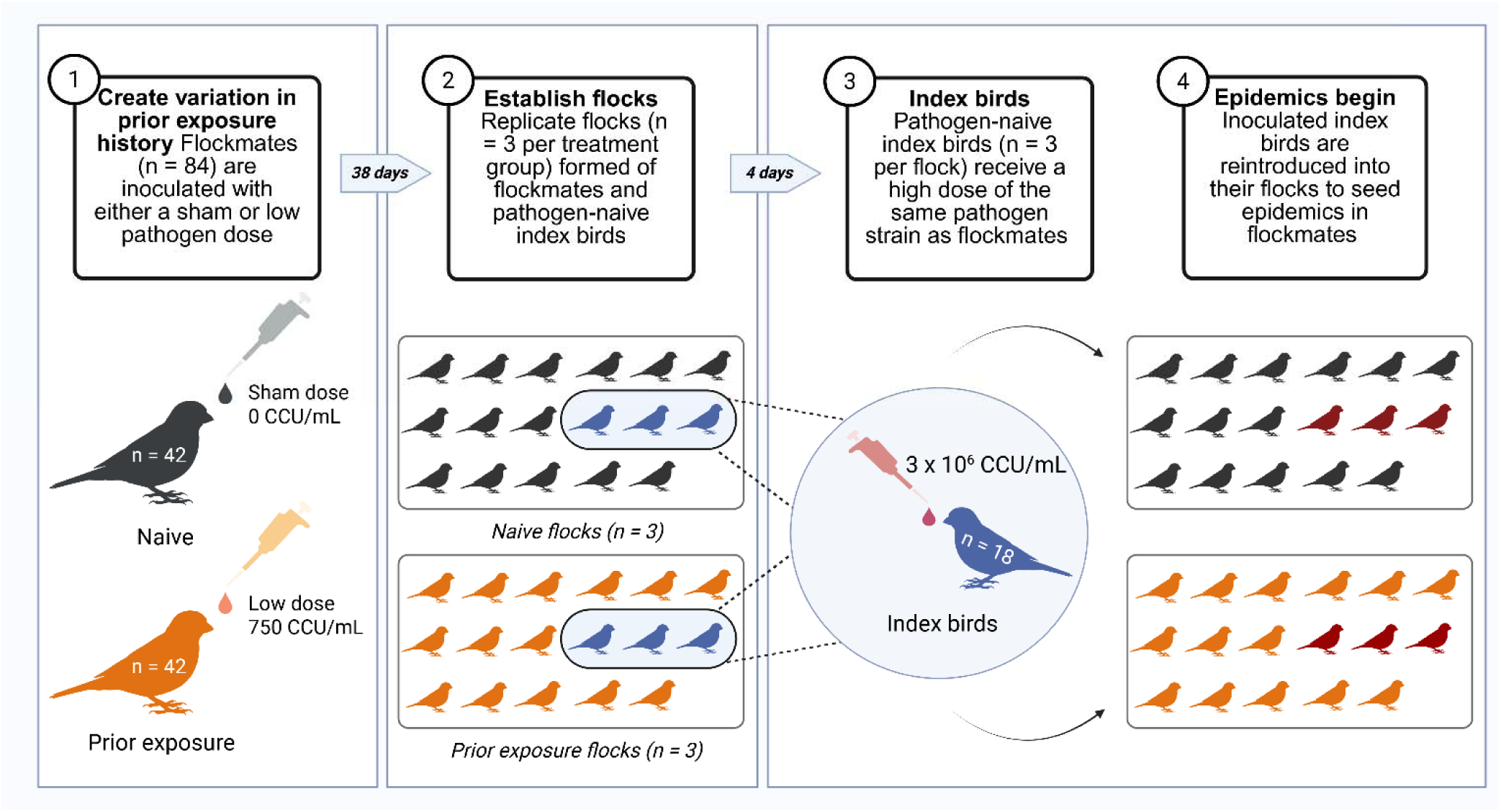
Experimental design. Flockmates (n = 84) were evenly split into two treatment groups: naïve (grey) vs prior exposure (orange). [1] First, flockmates received either a sham dose of sterile media (naïve birds) or a low dose of *M. gallisepticum* (prior exposure). Birds were allowed to recover from their primary exposure for more than a month. [2] Flocks of 17 birds, comprising 14 flockmates from the same treatment group and three initially pathogen-naïve index birds (blue), were established in six identical aviary units. [3] Index birds received a high inoculation dose to [4] seed epidemics among the flockmates. Arrows between panels indicate how many days passed between parts of the experiment. Created in BioRender. Perez-Umphrey, A. (2026) https://BioRender.com/5gfelrv.

**Table 1.** Summary of results by flock. Flocks (a - f) are divided by treatment group (naïve vs prior exposure flocks). For the prior exposure flocks, the number of birds (prevalence and count) that became infected during primary inoculation are shown. The sex ratio is male (M) to female (F). Duration of infection is listed in ascending order for each flock’s index birds and mean infection duration per flock is reported for infected flockmates. The mean of infected flockmates’ maximum pathogen load and eyescore are the log_10_ copies and total categorical value, respectively. Prevalence of infection and pathology (conjunctivitis) is the proportion of flockmates that ever exhibited that characteristic, reported as the percentage and count. Standard errors are given with mean values.

| Treatment group | Aviary flock | Sex ratio | Primary exposure infection prevalence % (n) | Duration of infection (days) |  | Infected flockmates |  | Flockmate prevalence |  |
| --- | --- | --- | --- | --- | --- | --- | --- | --- | --- |
| | | | | Index birds (in ascending order) | Flockmates (infected only) mean ± SE | Mean maximum pathogen load ($\log_{10}$ copy number) ± SE | Mean maximum eyescore ± SE | Infection % (n) | Pathology % (n) |
| Naïve | a | 9M:8F | -- | 14, 14, 21 | -- | -- | -- | 0.0 (0) | 0.0 (0) |
|  | c | 9M:8F | -- | 14, 14, 21 | -- | -- | -- | 0.0 (0) | 0.0 (0) |
| | e | 8M:9F | -- | 6, 14, 21 | 12.7 ± 4.3 | 4.84 ± 0.1 | 2.8 ± 1.0 | 21.4 (3) | 21.4 (3) |
| Prior Exposure | b | 9M:8F | 28.6 (4) | 6, 14, 35 | 5.0 ± 2.7 | 3.3 ± 0.8 | 2.0 ± 0.4 | 21.4 (3) | 14.3 (2) |
| | d | 12M:5F | 14.3 (2) | 14, 14, 35 | 10.1 ± 5.1 | 2.7 ± 0.5 | 1.5 ± 0.4 | 71.4 (10) | 28.6 (4) |
| | f | 8M:9F | 28.6 (4) | 6, 21, 21 | 1.0 ± 0.0 | 1.5 ± 0.1 | -- | 14.3 (2) | 0.0 (0) |

### Experimental timeline and sample collection

On day 0 of the experiment (i.e., the day primary inoculations took place; September 2^nd^, 2022), flockmates in the prior exposure group received a bilateral ocular inoculation of 70 µl (35 µl per eye) of 7.5 x 10^2^ color-changing units (CCU)/mL of MG suspended in Frey’s media. The MG isolate used was the original index isolate, VA1994 (Ley, Berkhoff, and McLaren 1996). The birds in the naïve group received a sham inoculation of the same amount of sterile Frey’s media. Thirty-eight days post inoculation (DPI), flocks were established by moving birds into each of the six aviary units (Figure 1). All prior exposure treatment birds were confirmed to be recovered – i.e., absence of clinical signs – before flock establishment. Pathogen load data were collected from all birds to ensure they were uninfected (defined as < 15 MG copies per assay [Appendix S1:Section S3]) before epidemics were initiated.

To seed epidemics, four days after establishing the flocks at the aviaries (October 14^th^; DPI 42), the three index birds in each flock received a high inoculation dose (3.0 x 10^6^ CCU/mL) of 50 µl (25 µl per eye) of the same MG isolate that the prior exposure flockmates had received. All index birds were successfully infected and presented clinical signs in one or both eyes by the first sampling occasion six days later (Appendix S1:Figure S1). On sampling days, birds were caught with butterfly nets and epidemic progression was monitored using eye swab (pathogen load) and eye score (pathology) data collected longitudinally throughout the experiment for each bird (Appendix S1:Table S2).

Disease severity was measured using a categorical scoring system where the severity of mycoplasmal conjunctivitis is scored from 0 - 3 in 0.5 increments for each eye, where 0 indicates there is no pathology and 3 is the most severe, for a total summed score of 0 - 6 per bird at a given time point (Sydenstricker et al. 2005). To collect pathogen load data, both conjunctiva were swabbed with small, sterile cotton swabs that were then vigorously dipped and swirled in 300 µl tryptose phosphate broth (TPB). The swabs were discarded and the TPB sample solution was frozen at −20°C until nucleic acid extraction and measurement via qPCR assay (Appendix S1:Section S4). The same two researchers (APU and DMH) blind to treatment scored and swabbed the birds throughout the experiment to minimize observer biases.

### Data Analysis

#### Statistical analysis

To understand the effects of prior exposure and associated heterogeneity in susceptibility on flock-level prevalence of infection, we fit a generalized linear mixed effects model (GLMM; *glmmTMB* (Brooks et al. 2017); R v. 4.3.1 [team 2024]) with a binomial distribution and a logit link. The response variable was individual presence/absence of infection (with infection defined as >15 copies of MG detected via qPCR) each day. Mean duration of index bird infection per flock and flock type were interacting fixed effects. Random effects were individual bird nested within flock to account for repeated measures (the sampling of the same birds and flocks over time). Model selection was performed using Akaike’s Information Criterion values adjusted for small sample sizes (Burnham and Anderson 2002; AIC_C_). Because we expected prior exposure to confer some degree of protection to re-infection and the development of conjunctivitis, we also modeled prevalence as defined by the absence (0) or presence (1) of pathology (> 0 eyescore) in the same manner. To account for the one flock with a skewed sex ratio, we asked whether an individual’s infection probability differed by sex in a GLMM with a binomial distribution and logit link, and where the response variable was 0 (uninfected) or 1 (ever infected; results presented in Appendix S1:Section S2 and Table S3). In models that included effects of flock type and mean infection duration of the index bird, we calculated variance inflation factors (VIF) using the package *car* (Fox and Weisberg 2019) to assess potential multicollinearity in the predictor variables. We centered the data (subtracted the mean value from all input values of the continuous predictor variable, such that the centered mean was zero; Schielzeth 2010) and re-ran the models to confirm that model parameter estimate stability (Schielzeth 2010) had improved. Where applicable, values reported in the results section are based on these adjusted models.

To ask whether prior exposure reduced infection or disease severity, we compared maximum pathogen loads and pathology by flock type. We limited this analysis to infected birds only, to ensure that the large number of flockmates that did not get infected – including all of the flockmates from two naïve flocks (see Results) – did not bias our results. Pathogen loads and pathology of infected birds did not follow a Gaussian or other described distribution, so we used nonparametric Wilcoxon rank sum tests.

#### Population model fitting

To determine whether prior exposure induced heterogeneity in susceptibility in the experimental epidemics, and whether, under more natural conditions, the observed heterogeneity in infection risk followed previous patterns of experimental dose-response data where exposure was controlled for each individual, we modeled epidemics within flocks using a continuous-time Markov chain SIR model.

Continuous heterogeneity in susceptibility was described using a gamma distribution that was truncated and discretized for simulations (described below; see Appendix S1:Section S5.1-5.3 equations for the equivalent deterministic model). Incorporating heterogeneity in susceptibility meant that the force of infection depended on an individual’s susceptibility, denoted by *x,* as well as on the transmission rate β (Table 2). This was compared to the equivalent homogeneous SIR model in which there is no variation in susceptibility, represented by a single class of susceptible individuals (see Appendix S1:Section S5.4-S5.6 equations). To allow for more direct comparisons between the heterogeneous and homogeneous models, we rescaled transmission rate between the two models by adding a parameter *a*, which summarizes the mean susceptibility between the two models where *a =1/k*θ and k and θ are shape and scale parameters, respectively, from the gamma distribution of susceptibilities in the heterogeneous model.

**Table 2.**
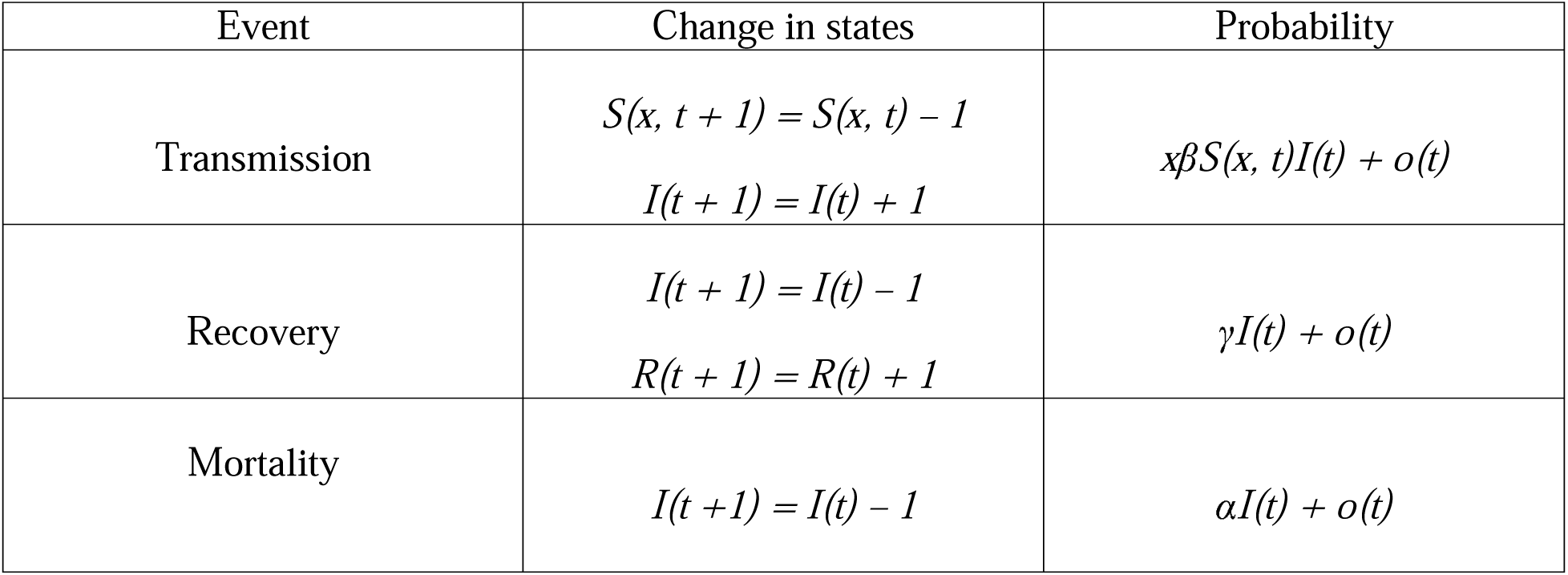
Stochastic population model events and probabilities. Initial conditions: *S(x,0) ∼Multinomial (n=S(0), p=Gamma(x))*

The use of a stochastic rather than a deterministic model allowed us to account for stochastic population processes that are relevant at small experimental population sizes (i.e., n = 17 birds per flock). Infection, recovery, and mortality were all treated as Markov processes which depend on the transition probabilities at time *t* but not on the history of previous events (Table 2). Transmission rate β and recovery rate γ were set at values derived from previous work in this system (Williams et al. 2014). Because transmission in this model depended on transmission rate β multiplied by host susceptibility x (or *a* in the equivalent homogenous model; see above and Appendix S1: Section S5.1-5.5 equations), we were able to estimate changes in transmission as changes in susceptibility x, even though transmission rate β was fixed. Mortality rate α was set at 0 for model fitting, under the assumption that disease-induced mortality is zero in captivity (Kollias et al. 2004).

Stochastic models were simulated using the Gillespie algorithm (Gillespie 1977) in R, which approximates the time to the next event τ as an exponential, and the probabilities of each event based on the states at time *t* and parameter values (Table 2). At the beginning of the simulation, each of the initially susceptible birds (n = 14) was randomly assigned a susceptibility *x* according to the gamma distribution *f(x)*. To discretize this continuous distribution, we truncated individual susceptibilities at 10 and divided the 0 to 10 range into 1000 evenly sized bins, chosen to match the discretization used in numerical simulations of the equivalent deterministic models in previous work (Hawley et al. 2024). With 1000 bins over the range 0 to 10, the mean and variance of susceptibility within the initial population of infected birds closely matched that of the continuous gamma distribution on average across 1000 model realizations. At the start of the epidemic, individuals were assigned to a susceptibility value according to a multinomial distribution (Appendix S1:Section S5), where the probability of each susceptibility bin was set by the gamma probability distribution function in R (*dgamma*). Therefore, heterogeneity in susceptibility became an additional source of stochasticity in the model: as the heterogeneity described by the gamma distribution increased, the probability of choosing fourteen individuals with high variance in susceptibility increased.

#### Model fitting

To ask how heterogeneity in susceptibility changed with prior exposure, we compared model fits to the experimental epidemic dataset using two candidate models fit to the data separately:

1) The heterogeneous SIR model described above, with a gamma distribution of susceptibilities within each flock.
2) A homogeneous SIR model assuming all individuals in a flock have equal susceptibility to infection.

For each of these models, we also compared two sets of parameter values for susceptibility:

1) Estimated from a previous study using experimental dose response data (Hawley et al. 2024).
2) Fitted directly to the experimental epidemic dataset using a Bayesian approach (outlined below).

By comparing these parameters sets, we were able to determine whether prior exposure-induced heterogeneity in host risk of infection (i.e., susceptibility) in a more naturalistic setting followed patterns of heterogeneity in susceptibility in experimentally controlled laboratory infections, where susceptibility was defined more narrowly as the likelihood of infection to a controlled exposure. For the heterogenous model, we estimated the shape and scale parameters for the gamma distribution of susceptibilities directly from the epidemic dataset for prior exposure and naïve flocks using an Approximate Bayesian Criteria (ABC) approach (Minter and Retkute 2019; described below). Models were fit to the prior exposure group and naïve groups separately (Table 3). We included data from all three flocks per treatment when fitting model parameters, including the two of three naïve flocks that had no evidence of ongoing transmission from the index birds (see Results). The absence of epidemics in some experimental populations is not unexpected given their small population sizes and the inherently high degree of stochasticity in these processes (see simulation Results). However, to ensure that our results were robust to the inclusion of the two naïve flocks where no epidemics occurred, we also compared model fits for the single naïve flock that had transmission (Appendix S1:Table S4).

**Table 3.** Model fitting for stochastic SIR. Better fitting models have a lower median and minimum discrepancy, measured as the sums of squares difference (residual sums of squares; RSS) between model realizations and the three (per treatment) empirical flock trajectories of infected birds over time. Parameters from dose response data were from a previously published study (Hawley et al. 2024) and experimental epidemic data were collected in the current study. Note that a coefficient of variation (CV) for susceptibility is not reported for homogenous models because these models do not allow for any variation in susceptibility. Sixty-six percent credible intervals are reported for mean susceptibility and CV. ^A^ Reported values are the value of mean susceptibility (and CV) from fitted dose response data parameters, which is not identical to the median of the bootstrapped confidence intervals. ^B^ Parameter for the homogenous dose response data model restricted between 0 and 1, therefore not directly comparable to the mean susceptibility of the heterogeneous model. ^C^ Dose response data confidence intervals are computed via 1000 bootstrapped samples with replacement from data, therefore not directly comparable to the credible intervals computed in this study.

| Treatment | Model | Number of parameters fitted (n) | Dataset for susceptibility parameter estimates | Mean susceptibility: median estimate (66% credible intervals) | CV susceptibility: median estimate (66% credible intervals) | Median Discrepancy (RSS) between model and flock data | Minimum Discrepancy (RSS) between model and flock data |
| --- | --- | --- | --- | --- | --- | --- | --- |
| Naïve | Homogeneous susceptibility | 1 | Dose response data (Hawley et al. 2024) | 0.838<sup>A,B</sup><br>(0.708, 1.000)<sup>C</sup> | -- | 1060.8 | 98.2 |
|  |  |  | Flock data (this study) | 0.062<br>(0.033, 0.10) | -- | 167.4 | 52.0 |
|  | Heterogeneous susceptibility | 2 | Dose response data (Hawley et al. 2024) | 1.18<sup>A</sup><br>(1.039, 1.530)<sup>C</sup> | 0.899<sup>A</sup><br>(0.805, 0.961)<sup>C</sup> | 217.9 | 32.9 |
|  |  |  | Flock data (this study) | 0.06<br>(0.022, 0.088) | 1.79<br>(1.28, 2.44) | 75.0 | 22.0 |
| Prior exposure | Homogeneous susceptibility | 1 | Dose response data (Hawley et al. 2024) | 0.140<sup>A,B</sup><br>(0.107, 0.246)<sup>C</sup> | -- | 164.1 | 41.0 |
|  |  |  | Flock data (this study) | 0.023<br>(0.0025, 0.12) | -- | 167.4 | 52.0 |
|  | Heterogeneous susceptibility | 2 | Dose response data (Hawley et al. 2024) | 0.446<sup>A</sup><br>(0.295, 0.687)<sup>C</sup> | 1.630<sup>A</sup><br>(1.230, 1.786)<sup>C</sup> | 174.0 | 36.0 |
|  |  |  | Flock data (this study) | 0.065<br>(0.013, 0.24) | 3.53<br>(1.98, 7.12) | 173.4 | 47.1 |

In the ABC approach, each set of parameters proposed from the prior distribution was used to simulate the model, which was then compared to the data using a summary statistic to compute the discrepancy between the model simulation and the data across three realizations. Parameter sets whose discrepancy from the data fell below an assigned tolerance threshold were retained in the posterior. Because we had three different experimental flock trajectories per treatment, we simulated three realizations of the model and compared each of these to one of the empirical infection trajectories (randomly matched). We used the sums of squares difference of the infected population trajectories at each empirically sampled time point to measure the discrepancy between the data and the model for each realization (Smith et al. 2005). We then summed that squared difference across all time points and all three experimental flocks to form our summary statistic.

While ABC algorithms often simulate only one realization per parameter set for computational efficiency, we instead chose to simulate 50x per parameter set (with three realizations each time) and recorded the median sums of squares discrepancy out of those 50 values. The use of more realizations (150 total) of the stochastic model to approximate the sums of squares discrepancy from the data was particularly valuable here due to the increased stochasticity in the initial conditions that was introduced by the discretized gamma distribution of heterogeneity in susceptibility in the heterogeneous model. The priors were vague lognormal distributions (σ^2^ = 2) centered on the gamma shape and scale values estimated from the previous dose response experiment, in which naïve and previously exposed birds were infected with the same MG strain as the current study (e.g., see Results, Table 3).

We used the same general ABC method to fit the homogeneous stochastic model to the prior exposure and naïve flock datasets. In the homogenous model, only a single parameter was fit (the mean susceptibility value), and there was only one S class (in other words, a traditional stochastic SIR model; see Appendix S1:Section S5.4-S5.6 for equivalent deterministic model equations). Finally, to compare goodness of fits of the parameters estimated from dose response data versus experimental epidemic data, we simulated the stochastic models 1000 x 3 times for both sets of parameter values (previous estimates from the dose response model and fitted estimates) for the heterogeneous and homogenous models (four total parameter sets). We then saved the median and minimum sums of squares discrepancy out of the 1000 realizations for each parameter set. Lower values of sums of squares discrepancy between the data and the model indicate a better fit to the data (Smith et al. 2005).

## Results

There was substantial variation in prevalence among flocks (0 – 71.4%; Figure 2; Table 1), with 2/3 experimental flocks in the naïve treatment showing no detectable transmission from index birds. Although we used identically-treated and all immunologically-naïve index birds (three index birds per flock) to seed the flock epidemics, we found unexpected variation in index bird competency (defined as pathogen transmission ability [Cortez and Duffy 2021; Gervasi et al. 2015]) across flocks and treatment groups, measured as both the average pathogen loads and lengths of infection for index birds. By chance, the longest index bird infections occurred in two of three prior exposure flocks (b and d) which each contained an index bird that did not resolve their infection during the period they were sampled (Figure 3; Table 1). Therefore, we accounted for this random variation among index birds in infection duration (6-35 days; Figure 3) by including mean index bird duration in statistical models of prevalence.

**Figure 2.**
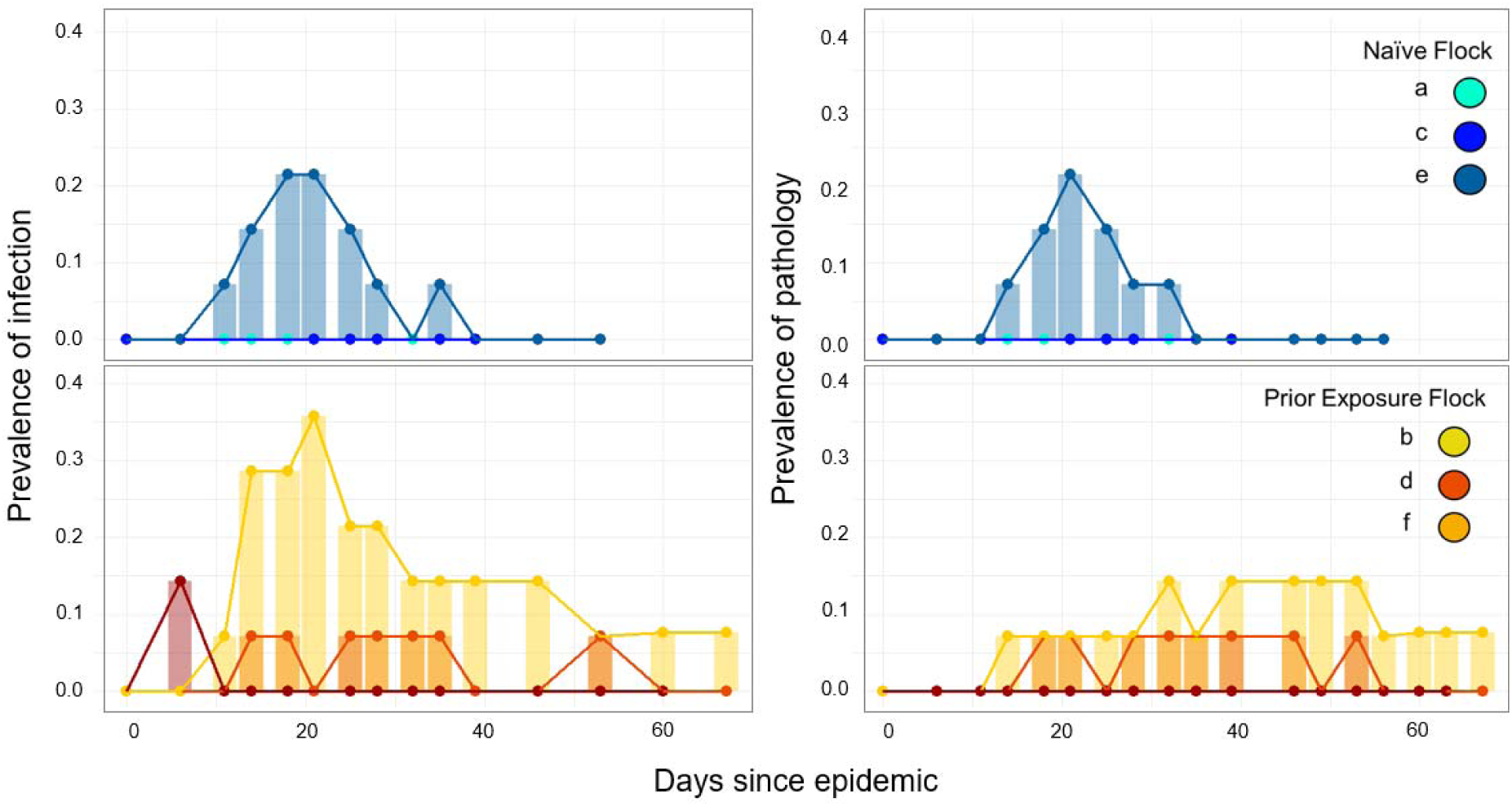
Prevalence of infection (left panels) and pathology (right panels) in flockmates (y-axes) over time since the inoculation of the index birds (not shown) in each flock (x-axis, t = 0 marks the start of the epidemic). Naïve (top panels) and prior exposure flocks (bottom panels) are distinguished by color.

**Figure 3.**
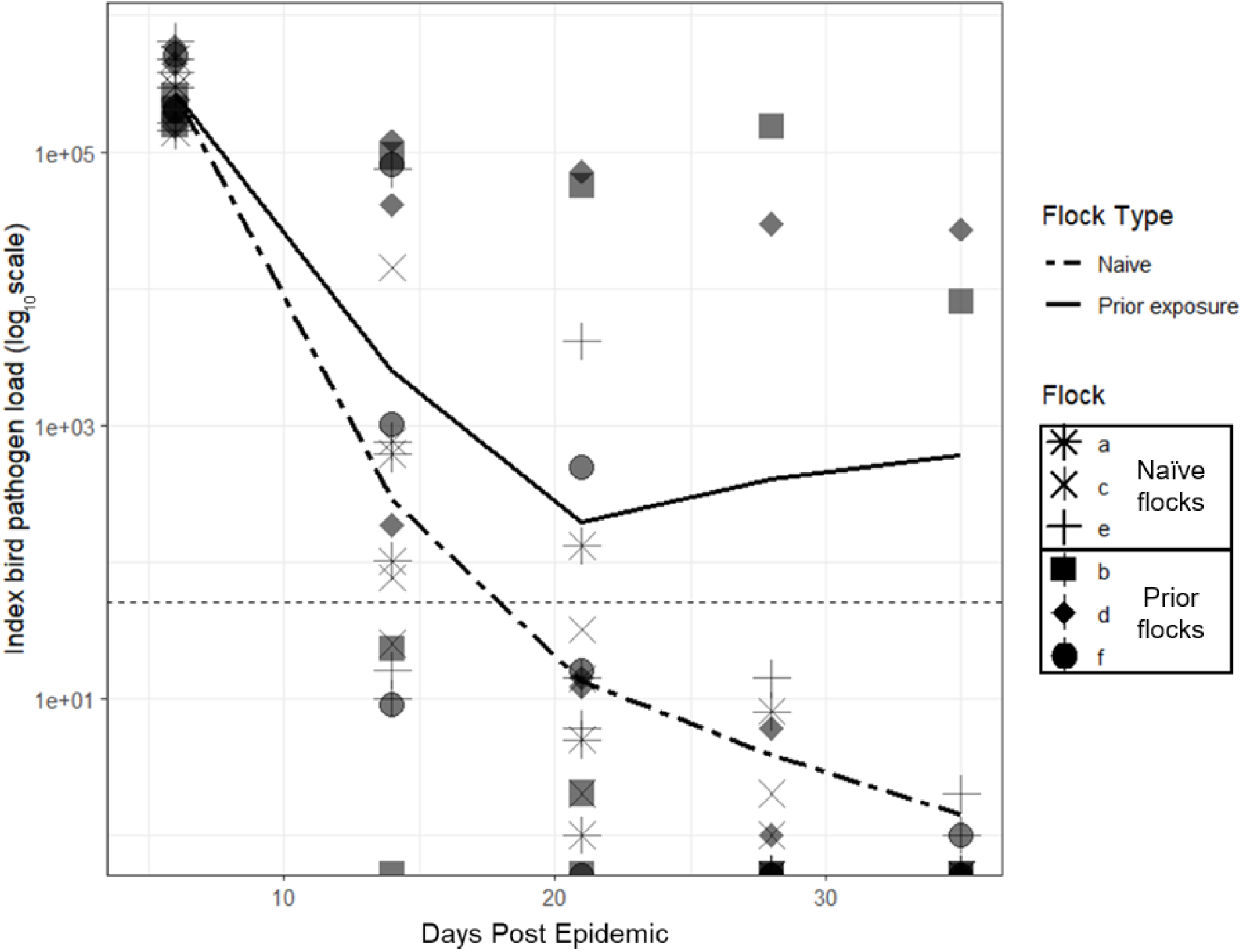
Pathogen loads (y-axis, log_10_ scale) per individual index bird over time since inoculation (x-axis, t = 0 marks the start of the epidemic). Flock is represented by the shape of the data points and lines show the mean per flock type (solid, prior exposure; dashed, naïve). The horizontal dashed line indicates the pathogen load cutoff used to determine infection status. Note that index birds in both treatments were pathogen-naïve when inoculated; differences between flock type are due to random chance and not to applied treatments.

Model selection using AICc identified the most informative model – a generalized linear mixed model with binomially distributed data and a nested random effect of bird within flock – as having an interaction between flock type and mean duration of index bird infection (prior exposure:mean index infection duration: coefficient ± 1 SE: 1.23 ± 0.544). A simpler model including only the fixed effect of mean index bird infection duration (coefficient ± 1 SE: 0.323 ± 0.156) received similar support (ΔAIC_C_ < 2; Appendix S1:Table S3). While average prevalence was higher in the prior exposure versus naïve flocks (Figure 2, Table 1), models that considered flock type as the sole predictor were not supported.

When prevalence was defined by the presence/absence of pathology, the top model was the null model (generalized linear mixed model with binomially distributed data and a nested random effect of bird/flock, including only the intercept; β_0_ = −7.99 ± 1.05), indicating that flock type did not improve model explanatory power of the prevalence of pathology. Models with single fixed effects (mean index infection duration: coefficient ± 1 SE = 0.074 ± 0.233; prior exposure: coefficient ± 1 SE = 0.167 ± 1.044) ranked similarly to the top model (ΔAIC_C_ < 2; Appendix S1:Table S3).

On an individual level, prior exposure appeared to induce protection in birds that became detectably infected (qPCR+) during the epidemics, particularly from disease (Figure 4). In Wilcoxon rank sum tests, infected bird maximum pathogen load did not significantly differ by flock type (W = 39, n = 3 naïve, 15 prior exposure, p = 0.0564). However, maximum pathology was greater in naïve flockmates (W = 41.5, n = 3 naïve, 15 prior exposure, p = 0.0184; [median] prior exposure flockmates = eyescore 0, [median] naïve flockmates = eyescore 3), suggesting that prior pathogen exposure protects birds from severe disease upon reinfection, even if pathogen loads do not vary as strongly. Whereas all infected naïve birds also had detectable conjunctivitis, in prior exposure flocks, most infected birds were aclinical. This resulted in discrepancies between the prevalence of infection versus pathology, where the total number of infected birds (n = 15) exceeded that with conjunctivitis (n = 9; Figure 2 left versus right panels; Table 1) in prior exposure flocks.

**Figure 4.**
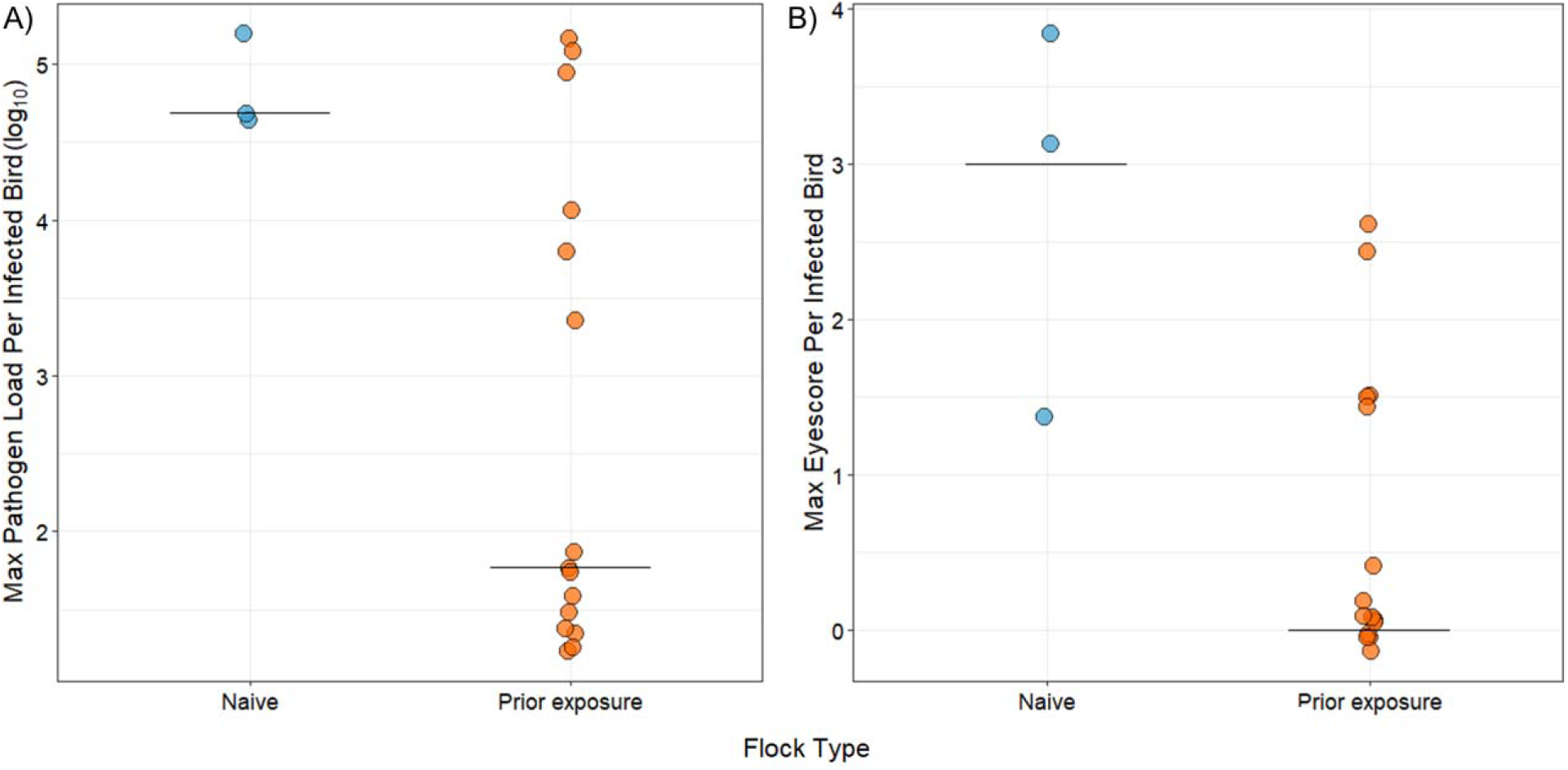
Flockmates with prior exposure (flock type) showed moderately, but not significantly, lower maximum pathogen loads (A), and significantly lower maximum eyescores (B) relative to flockmates that were pathogen-naïve when successfully infected during the epidemic. The line indicates group median and is overlaid by datapoints of maximum pathogen load (log_10_ scale; left) and eyescore (right) for flockmates ever infected during the epidemics (> 15 pathogen copies). Naïve flockmates are on the left of each panel (blue) and prior exposure flockmates are on the right (orange).

### Population model fitting

When stochastic models were fit to the data from each treatment (Figure 5), heterogeneity in susceptibility was greater for prior exposure flocks (CV [66% credible intervals] = 3.53 [1.98, 7.12]) relative to naïve flocks (CV = 1.79 [1.28, 2.44]) when modeled with gamma-distributed heterogeneity in susceptibility (Table 3). However, the estimated CV of 1.79 for pathogen-naïve flocks, while lower than prior exposure flocks, was higher relative to heterogeneity estimates for pathogen-naïve birds from prior dose response experiments in individual cages (CV = 0.899 for pathogen-naïve birds in Hawley et al. 2024; Table 3). Using the epidemic data from naïve flocks, models with heterogeneity in susceptibility were preferred to homogenous models (Table 3), providing further support for notable heterogeneity in susceptibility even in pathogen-naïve flocks in this study.

**Figure 5.**
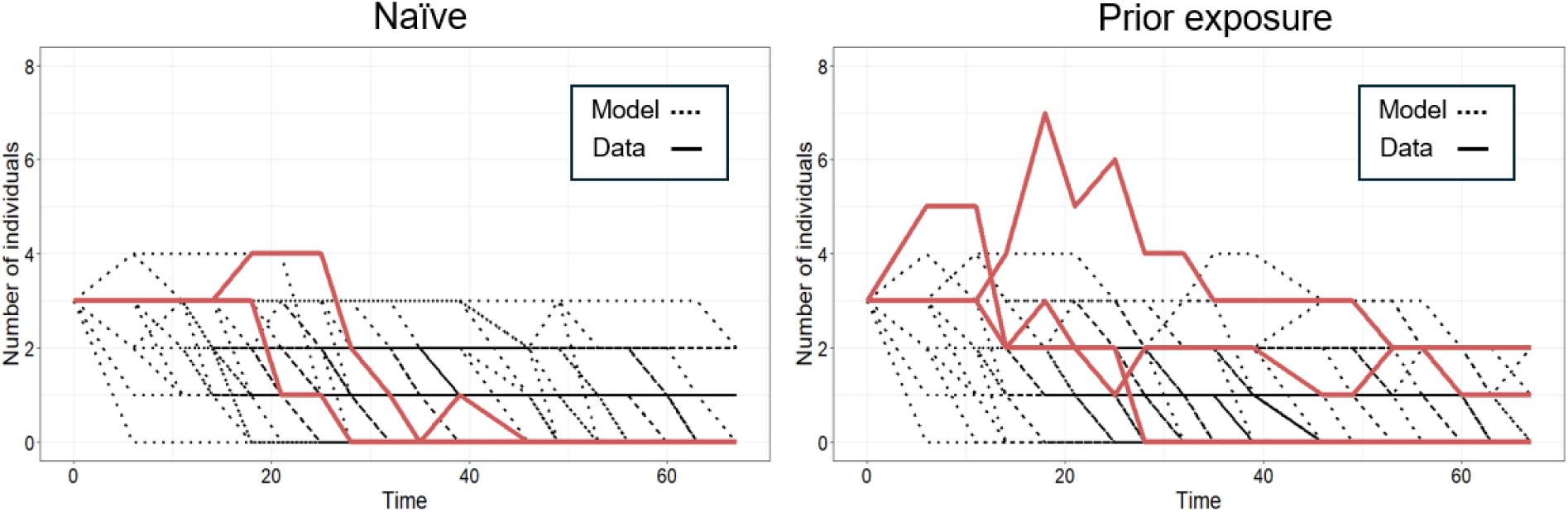
Stochastic model realizations (black dotted lines; n = 100 realizations) and empirical data (red solid lines) for the number of infected individuals from an SIR model with heterogeneity in susceptibility, using best-fitting parameters for experimental flocks that were pathogen-naïve (left panel) or had prior exposure (right). Only the infected class, to which the models were fit, is shown [see Appendix S1:Figure S2 for all classes]). Heterogeneity was described as a gamma distribution (naïve flocks: k = 0.312, θ = 0.129; prior exposure flocks: k = 0.08, θ = 0.783). Transmission (β = 0.00275) and recovery rates (γ = 0.03) were from prior literature (Hawley et al. 2024; Williams et al. 2014). Mortality α was set to zero because infection-induced mortality did not occur in captivity.

When we fit the stochastic model parameters to the data from the single naïve flock with detectable transmission, estimates for heterogeneity in susceptibility remained similar (CV = 1.59 [1.09, 2.29]) to that estimated for all three naïve flocks combined (CV = 1.79 [1.28, 2.44]), with largely overlapping credible intervals (Appendix S1:Table S4). For models fit to only the single naïve flock, both the heterogeneous and homogenous models yielded slightly higher estimates for mean susceptibility than when all naïve flocks were analyzed together (Appendix S1:Table S4). However, restricting the analysis to this single naïve flock did not improve model fits (Appendix S1:Table S4).

While prior exposure flocks showed a similar qualitative pattern of increased estimates of heterogeneity (CV = 3.53) compared to those derived from published dose response experiments in individual cages (CV = 1.630 in Hawley et al. 2024 for birds with prior exposure; Table 3), all four stochastic models considered (i.e., homogeneous and heterogeneous with parameter values from dose response versus fit to the flock data) produced similar fits using the residual sums of squares values (median or minimum across 100 model realizations) to compare predicted versus observed numbers of infected birds in prior exposure flocks (Table 3). While the median discrepancy across 100 realizations was slightly lower for the homogeneous model than the heterogeneous model, the opposite was true for the minimum discrepancy across those same realizations. This indicates that, while more of the simulated trajectories from the homogeneous model were slightly closer to the observed data in the prior exposure flocks, the best matching individual realization for a given parameter set was slightly better for the heterogeneous model.

## Discussion

Infectious disease outbreaks and their characteristics are fundamentally driven by pathogen and host traits (Gomes et al. 2022; Lloyd-Smith et al. 2005; VanderWaal and Ezenwa 2016), which can vary substantially in natural populations (Vazquez-Prokopec et al. 2016). Although continuous distributions of host traits are increasingly included in epidemiological models (Gomes et al. 2022; Tuschoff and Kennedy 2024; Corder, Ferreira, and Gomes 2020), experimental transmission studies that integrate theoretical predictions and empirical data remain rare. Here, we tested whether prior-exposure induced heterogeneity in susceptibility, and resulting experimental epidemics, aligned with theoretical predictions generated from a previous dose-response experiment in the house finch-*M. gallisepticum* system (Hawley et al., 2024). Consistent with previous work (Hawley et al., 2024), heterogeneity in susceptibility was estimated as being higher in flocks with prior pathogen exposure relative to pathogen-naïve flocks. However, despite the heterogeneous model performing slightly better, prior exposure flocks were described equally well by homogenous as heterogeneous models, and so there was no “best fit”. A key difference from past findings was that, while pathogen-naïve birds in the dose response study (performed using individually-housed birds) were best described by models that assume homogenous susceptibility, pathogen-naïve flocks in our experimental epidemics were best described by models that incorporated gamma-distributed heterogeneity (although, we note that the differences in parameter numbers across studies limits this comparison). This result indicates that in a more “natural” mesocosm setting, infection risk can be heterogenous even for a population with no previous pathogen exposure. It appears that the flock epidemics captured additional sources of heterogeneity, such as variable host competence, that are absent from more controlled experiments.

Contrary to theoretical predictions (Langwig et al. 2017; Ben-Ami, Ebert, and Regoes 2010; Katriel 2012; Gomes et al. 2014; Gomes et al. 2022; Dwyer, Elkinton, and Buonaccorsi 1997; Hawley et al. 2024) and our prior work (Hawley et al. 2024), flocks with prior exposure and greater estimated heterogeneity in susceptibility did not have lower mean prevalence during experimental epidemics. However, the lack of transmission in two of the three naïve flocks makes it difficult to determine whether this was due to treatment or a lack of exposure opportunities. In fact, we found that among-flock prevalence was significantly predicted by the duration of index bird infection alone and in interaction with flockmate treatment, despite our attempts to standardize index bird infectiousness by using all naïve index birds in our experimental design (for both naïve and prior-exposed flockmate treatments). The index birds, which were all inoculated with the same high pathogen dose, varied substantially in their time to pathogen clearance, which, conservatively, ranged from 6 to 35 days (Figure 3; Appendix S1:Figure S1). Moreover, the random placement of the two index birds with the longest infections in two of the three prior exposure flocks appeared to be the strongest driver of flockmate infection and disease prevalence in our experiment. The presence of individual hosts with disproportionately high infectiousness (i.e., “superspreaders”) is a common and important feature of disease dynamics (Lloyd-Smith et al. 2005; Woolhouse et al. 1997; VanderWaal and Ezenwa 2016; Vazquez-Prokopec et al. 2016), and can cause less frequent but more intense (occurring faster and reaching higher epidemic peaks) disease outbreaks (Lloyd-Smith et al. 2005; Elie, Selinger, and Alizon 2022; Woolhouse et al. 1997). While individual variability in transmission is a product of multiple underlying factors (i.e., environmental conditions, immunological status, physiology, behavior [Stein 2011; Paull et a. 2012]), the index birds in this present study showed clear inter-individual variability in host competence – as measured by their infection duration – with which infectiousness, or pathogen shedding, was strongly correlated (Figure 3; Appendix S1: Figure S1).

An intriguing additional, and non-mutually exclusive, possibility for why prior exposure flocks did not have lower prevalence as predicted, is that birds in those flocks were more likely to be aclinical carriers (i.e., infected but with no pathology). Indeed, prior exposure provided protection to birds at the individual level, such that birds with prior exposure had significantly lower maximum pathology if infected during the epidemics, and most of the infected flockmates in prior exposure flocks remained entirely aclinical. This raises the possibility that the lack of pathology in infected birds with prior exposure allowed for better maintenance of normal behaviors during infection, which can be positively correlated with transmission in this system (Adelman and Hawley 2017; Ruden and Adelman 2021), and facilitated aclinical pathogen spread. On the other hand, more severe conjunctivitis is also known to promote transmission in this system (Hawley et al. 2023; Henschen et al., 2025), in which case the lower maximum pathology of infected flockmates would reduce transmission in prior exposure flocks relative to naïve flocks.

Disease epidemics, which typically begin with only a few infected individuals, are inherently stochastic and thus subject to die out (e.g., Lloyd-Smith et al. 2005). Indeed, our stochastic simulations revealed highly variable outcomes for these small epidemics, highlighting the challenge of capturing transmission dynamics in a naturalistic setting. To capture the full range of stochastic outcomes, we included all flocks in our parameter estimation and simulations, despite the absence of ongoing transmission in two of our six flocks. While this makes it challenging to estimate the relative importance of prior exposure for epidemic outcomes, the exclusion of no-transmission flocks could bias our results and hide important biological phenomena, such as the differences in host competence among index birds that we observed. When we fit the stochastic model parameters to the data from the single naïve flock with transmission, we obtained slightly higher estimates for mean susceptibility than when all naïve flocks were analyzed together, as would be expected given that flocks without transmission reduce estimates of mean susceptibility. While these susceptibility estimates from the single flock were more consistent with previous work showing that prior exposure reduces mean susceptibility (Hawley et al. 2024), the differences were minor and restricting the analysis to this single naïve flock did not improve model fits. Further, estimates of heterogeneity in susceptibility, the main parameter of interest in our study, from the single naïve flock were statistically indistinguishable from estimates made using all three naïve flocks. Overall, the global increase in heterogeneity for both flock types under mesocosm conditions may have masked our ability to detect any potential dampening effects of heterogeneity in susceptibility between the treatments. Additionally, it is possible that the experimental epidemics were simply too short to fully capture the epidemiological impacts of heterogeneity in susceptibility. Our stochastic simulated epidemics had not yet burned out by the timepoint at which they were truncated (67 days) to correspond with the experimental epidemic data. Among the many challenges of quantifying heterogeneity in susceptibility empirically, is reliably measuring it during the early stages of an outbreak and at small sample sizes (Tuschoff and Kennedy 2024). These challenges are difficult to feasibly remedy for systems that use wild-caught vertebrates in captivity. However, recently developed methods that harness contact tracing data (Tuschoff and Kennedy 2024) could be used in the future to measure heterogeneity in susceptibility in the earlier stages of an epidemic, which, incidentally, may include the entire time span of this study.

The substantial differences in our estimates for mean susceptibility and heterogeneity in susceptibility (Table 3) between our flock mesocosm study and prior work on individually-housed birds (Hawley et al. 2024) are important but not surprising. The probability of infection, and thus mean susceptibility, is likely to be lower in a setting where transmission is natural (i.e., from other birds or fomites versus direct inoculation), the exposure dose is uncontrolled, animals can freely move and interact/avoid one another, and hosts and pathogen are exposed to ambient conditions. The discrepancy in the conclusions generated by controlled experiments (which estimate that a naïve population’s response is relatively homogenous for the two studies done to date [Hawley et al. 2024; Langwig et al. 2017] versus mesocosm epidemics, highlights the importance of quantifying heterogeneity in infection risk in more naturalistic settings. These results also give even greater credence to the necessity of incorporating heterogeneity in susceptibility in epidemiological models when fitting real-world data (Gomes et al. 2022). Not only does the incorporation provide more realistic epidemiological estimates, it also offers an opportunity to deconstruct the factors (e.g., behavioral, environmental, endogenous traits) with the greatest contributions to the observed dynamics.

In conclusion, we found that prior pathogen exposure contributed to the host heterogeneity in susceptibility that we observed in our experimental epidemics, offering further support for the importance of variable acquired immune protection (likely as a function of waning or incomplete immunity [Gomes et al. 2014; Le et al. 2021]) and its role in determining a population’s disease dynamics and subsequent host-pathogen evolution (Rodrigues et al. 2009; Rose et al. 2021; Hawley et al. 2024; Langwig et al. 2017; Fleming-Davies et al. 2015; Fleming-Davies et al. 2018; Read et al. 2015). Intriguingly, however, other sources of heterogeneity were present in the more naturalistic mesocosm setting, which may have masked any detectable epidemiological consequences of prior exposure-induced heterogeneity in susceptibility. Lloyd et al. (2020) argues that future work in this field should focus on “coupled” heterogeneities – where heterogeneity of a trait such as susceptibility or transmission extends beyond a single source of variation, but rather is modeled as a function of multiple covarying and coincident sources, such as coupled heterogeneities in infectiousness, contact rates, or infection duration (Paull et al. 2012; Vazquez-Prokopec et al. 2016). Broadly speaking, incorporating multiple types (i.e., susceptibility, transmission) of heterogeneity and multiple sources of that variation are key to understanding disease dynamics in human and wildlife populations.

## Acknowledgements

This research was made possible by grant R01GM144972 from the National Institutes of Health (NIH) Ecology and Evolution of Infectious Diseases (EEID) program. All birds were handled and captured under U.S. Fish and Wildlife Service (MB154804-0) and Virginia Department of Game and Inland Fisheries (066646) permits. All housing and experimental protocols were approved by the Virginia Tech Institutional Animal Care and Use Committee (IACUC). We thank Edan Tulman and Steve Geary (University of Connecticut) for providing the pathogen inoculum and plasmids, as well as Bill Hopkins (Virginia Tech) for facilitating our use of the Research Aviaries. We also thank Danielle Alms, Madeline Alt, Alicia Arneson, Annabel Coyle, Zhang Gao, Noelle Hodges, Sadoni James, Marissa Langager, Riley Meyers, Sara Teemer, and Caro Vela for assistance with experimental sampling.

## Notes

### Competing Interest Statement

The authors have declared no competing interest.

### Summary of Updates

The manuscript was revised based on reviewer comments. No overall conclusions have changed, however, important results supporting those conclusions have either been updated, removed, or been added to: Because of the unexpected lack of transmission in two of the flocks: 1) The analysis of infection loads over time for all flockmates was removed; and 2) The homogeneous and the heterogeneous stochastic models were re-fitted to data from naive flock (e) alone, excluding the two naive flocks where no epidemics occurred. This was done to show that the results are robust to the inclusion of the flocks with no transmission. Figure 4 was changed so that it is no longer a boxplot but instead shows group medians. Credible intervals equivalent to 1 standard error (i.e., 66% credible intervals) were added to the estimated parameters in Table 3. Values for mean and CV of susceptibility in Table 3 were updated. Figure 5 was remade so that it shows only the infected class of the SIR models.

