## Supplementary materials for "Infected host competence overshadows heterogeneity in susceptibility in shaping experimental epizootics"

**Appendix S1**

**Anna A. Pérez-Umphrey, Kate E. Langwig, James S. Adelman, Lauren M. Childs, Jesse Garrett-Larsen, Dana M. Hawley, and Arietta E. Fleming-Davies.** *Infected host competence overshadows heterogeneity in susceptibility in shaping experimental epizootics*

**Section S1: House finch capture, quarantine, and housing**

House finches were captured in Montgomery County, VA from the end of May to mid-August 2022. Birds were captured either using mist-nets or hand-built wire cage traps at feeders that were consistently filled with black oil sunflower seed to bait the birds. Birds were given captive metal bands and pair-housed in 46 x 76 x 46 cm cages in an indoor animal facility. Animals had access to food (20% sunflower seed and 80% pelleted diet: Roudybush Daily Maintenance Nibbles [Roudybush, Inc., Cameron Park, CA, USA]) and water *ad libitum*. Water was periodically treated with Bene-Bac Plus FOS and Probiotics (Pet-Ag, Hampshire, IL) and Endocox, an anticoccidial. Temperatures were kept between 20 – 26°C and lights were on a 12-hour light:dark cycle until birds were moved to flock-housing in an outdoor avairy (see below). Birds were sexed in late-August by plumage, at a time when pre-basic molt was still ongoing, resulting in some mis-sexing (see Experimental Design).

To experimentally manipulate host prior exposure, only immunologically naïve birds were used in this study. To this end, only hatch year birds – aged by plumage and the presence of yellow gapes – were captured and brought into the laboratory. To ensure that captured birds did not have prior exposure to MG, all birds were quarantined for 21 days and checked for conjunctivitis at scheduled intervals (days 3-5 and 8-11). Blood was collected from the brachial vein between the 14$^{th}$ - 21$^{st}$ day of quarantine for plasma samples which were used to check for the presence of antibodies in a *M. gallisepticum* antibody enzyme-linked immunosorbent assay (IDEXX, Westbrook, ME, USA), using a 0.061 OD (optical density) threshold (modified from Hawley et al. 2011). Seropositive birds or those that were housed with a bird that developed conjunctivitis during their quarantine were not used in this study.

For the controlled prior exposure treatment, birds were moved to individual housing within the same indoor facility where they were quarantined. Index birds (n = 18; 9 female and 9 male) were also individually housed under the same conditions but kept in a separate indoor facility. All birds received metal captive bands. Additionally, index birds received a single plastic color band. For the experimental epidemics, flockmates and index birds were moved from their separate indoor facilities to Virginia Tech's Research Aviaries in the Wild Animal Research Facility. Flocks were established in six large replicate aviary units (2.4 x 2.4 x 5.5 m), which are partially enclosed but exposed to the environment so that animals experience ambient weather and light conditions. The aviary units were set up identically: each had four hanging water dishes, one large 6-port feeder filled with the same food mixture they received previously, 8 dowel-rod perches, and two heat lamps (on a timer overnight from 17:00-10:00 hours), and a 6-foot plastic Christmas tree. A secondary food source (a dish placed on the floor) was later added to ensure all birds had uninterrupted access to food.

### Section S2: Sex differences in infection likelihood

Infection likelihoods did not differ between the sexes (generalized linear mixed model with binomially distributed data with the random effect of flock) in the top model, which included an interaction between the fixed effects of sex and flock type (sex [male]: coefficient = -0.782 ± 1.32, $z = -0.593$, $p = 0.55$; flock type [prior]: 1.08 ± 1.50, $z = 0.720$, $p = 0.47$; sex [male] * flock type [prior]: coefficient = 2.34 ± 1.56, $z = 1.501$, $p = 0.13$).

The null model and model with only the fixed effect of sex ranked similarly ($\Delta AIC_C < 2$; Appendix S1: Table S3).

### Section S3: Defining infection

We defined a bird as being infected if the pathogen load in its swab was greater than 15 *mgc2* copies per 3µl qPCR reaction. This cutoff allowed for the removal of any low-level environmental contamination potentially detected in the swab samples. A bird was considered as having pathology if it had a total eyescore greater than or equal to one. This allowed for the removal of low eyescores attributable to injuries that present as conjunctival swelling. One naïve bird (a male from flock c) was conservatively removed from the analysis due to a positive swab sample (115 MG copies) collected on DPE 0, although there were no other indications throughout the experiment that this bird was ever infected, and this was likely a sample contaminated either during its collection or extraction/assay (Leon and Hawley 2017).

### Section S4: Pathogen load quantification

DNA extractions from eye swab samples were performed using 96-well DNeasy Blood and Tissue Kits (Qiagen, Hilden, Germany). Three negative controls were included per plate and samples were lysed overnight at 56°C. Pathogen loads were estimated via a probe-based qPCR assay that targets the *mgc2* gene in MG (Grodio et al. 2008; Hawley et al. 2011). Assays were prepared with a QuantiNova Kit (Qiagen, Hilden, Germany) and run on a QuantStudio 5 machine. Reaction volumes and cycling conditions were those specified by the manufacturer. Samples were run against a standard curve of 1:10 serial dilutions of $3.86 \times 10^1$ – $3.86 \times 10^8$ copies of a pCR4-TOPO plasmid with *a mgc2* insert. Standards were run in triplicate and three negative controls were included on each assay plate.

**Section S5: SIR models**

The deterministic version of the heterogeneous SIR model fit to flock epidemics is an ordinary differential equation model with gamma distributed susceptibility $x$ (eqs. Appendix S1: Section S5.1-S5.3). Note that the deterministic equations shown below were not used in this study. Rather, it is shown here to inform comparisons with the results of Hawley et al. (2024). Infection of an individual of susceptibility $x$ occurs at a scaled transmission rate $x\beta$. Once infected, all individuals move to a single I class, integrated over all possible susceptibility values from 0 to infinity. Infected individuals recover at rate γ. Disease induced mortality μ was set to zero for experimental epidemics as mortality in captivity is approximately zero although it is non-zero in natural settings.

(5.1) $$\frac{dS(x)}{dt} = -x\beta S(x)I$$

(5.2) $$\frac{dI}{dt} = \beta I \int xS(x)dx - \gamma I - \mu I$$

(5.3) $$\frac{dR}{dt} = \gamma I$$

<u>Equivalent homogenous SIR model</u>

(5.4) $$\frac{dI}{dt} = -\alpha\beta SI$$

(5.5) $$\frac{dI}{dt} = \alpha\beta SI - \gamma I - \mu I$$

(5.6) $$\frac{dR}{dt} = \gamma I$$

**Table S1.** Date (in the year 2022), associated epidemic sampling timepoint, and number of index birds and flockmates in each flock at that timepoint. All flocks began with n = 3 index (I) birds and n = 14 flockmates (F) with an approximately even sex ratio. Any decreases in sample sizes reflect incidental (i.e., not disease-related mortality). Beginning on the day a flock was removed from the study, sample sizes are 0.

| | | A | | B | | C | | D | | E | | F | |
|---|---|---|---|---|---|---|---|---|---|---|---|---|---|
| **DPE** | **Date** | **I** | **F** | **I** | **F** | **I** | **F** | **I** | **F** | **I** | **F** | **I** | **F** |
| 0 | 10/14 | 3 | 14 | 3 | 14 | 3 | 14 | 3 | 14 | 3 | 14 | 3 | 14 |
| 6 | 10/20 | 3 | 14 | 3 | 14 | 3 | 13 | 3 | 14 | 3 | 14 | 3 | 14 |
| 11 | 10/25 | 3 | 14 | 3 | 14 | 3 | 13 | 3 | 14 | 3 | 14 | 3 | 14 |
| 14 | 10/28 | 3 | 14 | 3 | 14 | 3 | 13 | 3 | 14 | 3 | 14 | 3 | 14 |
| 18 | 11/1 | 3 | 14 | 3 | 14 | 3 | 12 | 3 | 14 | 3 | 14 | 3 | 14 |
| 21 | 11/4 | 3 | 14 | 3 | 14 | 3 | 11 | 3 | 14 | 3 | 14 | 3 | 14 |
| 25 | 11/8 | 3 | 14 | 3 | 14 | 3 | 11 | 3 | 14 | 3 | 14 | 3 | 14 |
| 28 | 11/11 | 3 | 14 | 3 | 14 | 3 | 11 | 3 | 14 | 3 | 14 | 3 | 14 |
| 32 | 11/15 | 3 | 14 | 3 | 14 | 3 | 11 | 3 | 14 | 3 | 14 | 3 | 14 |
| 35 | 11/18 | 2 | 14 | 3 | 14 | 3 | 11 | 3 | 14 | 3 | 14 | 3 | 14 |
| 39 | 11/22 | 2 | 13 | 3 | 14 | 3 | 11 | 3 | 14 | 3 | 14 | 3 | 14 |
| 46 | 11/29 | 1 | 13 | 3 | 14 | 3 | 11 | 3 | 14 | 3 | 13 | 2 | 14 |
| 49 | 12/2 | 1 | 13 | 3 | 14 | 3 | 11 | 3 | 14 | 3 | 13 | 2 | 14 |
| 53 | 12/6 | 1 | 13 | 3 | 14 | 3 | 11 | 3 | 14 | 3 | 13 | 2 | 14 |
| 56 | 12/9 | 0 | 0 | 3 | 14 | 0 | 0 | 3 | 14 | 0 | 0 | 2 | 14 |
| 60 | 12/13 | 0 | 0 | 3 | 14 | 0 | 0 | 3 | 13 | 0 | 0 | 2 | 14 |
| 63 | 12/16 | 0 | 0 | 3 | 14 | 0 | 0 | 3 | 13 | 0 | 0 | 0 | 0 |
| 67 | 12/20 | 0 | 0 | 0 | 0 | 0 | 0 | 0 | 0 | 0 | 0 | 0 | 0 |

**Table S2. Experimental timeline.** Date and associated sampling timepoint (DPI: days post inoculation; DPE: days post epidemic), which are numbered in relation to day 0, or the inoculation and epidemic start days, respectively. All samples were collected in 2022. An "X" indicates whether a flock (a – f) was sampled at a given timepoint, what sample types were collected, and for which bird type (flockmate vs. index). Blank cells indicate that the sample was either not collected or unavailable (i.e., the flock had been removed from the study, or sample day/type was not relevant to that flock type or aviary role). On the first day of the epidemics (the day index birds were inoculated and re-released back into their flocks; days post epidemic [DPE] 0) and on DPE 6, all birds (flockmates and index birds) were swabbed and scored (Appendix S1:Table S1). Beginning on DPE 11, all birds were eye scored twice per week through the end of the experiment (DPE 67). Eye swabs were collected from flockmates twice a week between DPE 11 – DPE 39 and index birds' eyes were swabbed once per week during that period. After DPE 39, swabs were only collected from flockmates and only once per week. After DPE 49, flocks that had no evidence of pathology or transmission for three weeks were removed from the study before the final sampling day.

| **Time Points** | | **Flocks Sampled** | | | | | | **Samples Collected** | | | | | | | |
|---|---|---|---|---|---|---|---|---|---|---|---|---|---|---|---|
| | | **PRIOR** | | | **NO PRIOR** | | | **FLOCKMATES** | | | | **INDEX** | | | |
| **Date** | **Sample Timepoint** | **A** | **C** | **E** | **B** | **D** | **F** | **Swabs** | **Scores** | **Plasma** | **Inoculate** | **Swabs** | **Scores** | **Plasma** | **Inoculate** |
| 8/29 | **DPI -4** | X | X | X | X | X | X | X | X | X | | | | | |
| 9/2 | **DPI 0** | X | X | X | X | X | X | | | | X | | | | |
| 9/9 | **DPI 7** | X | X | X | X | X | X | X | X | | | | | | |
| 9/16 | **DPI 14** | X | X | X | X | X | X | | X | | | | | | |
| 9/23 | **DPI 21** | X | X | X | X | X | X | | X | | | | | | |
| 9/30 | **DPI 28** | X | X | X | X | X | X | | X | | | | | | |
| 10/7 | **DPI 35** | X | X | X | X | X | X | | X | X | | | | | |
| 10/14 | **DPE 0** | X | X | X | X | X | X | X | X | | | X | X | | X |
| 10/20 | **DPE 6** | X | X | X | X | X | X | X | X | | | X | X | | |
| 10/25 | **DPE 11** | X | X | X | X | X | X | X | X | | | | X | | |
| 10/28 | **DPE 14** | X | X | X | X | X | X | X | X | | | X | X | | |
| 11/1 | **DPE 18** | X | X | X | X | X | X | X | X | | | | X | | |
| 11/4 | **DPE 21** | X | X | X | X | X | X | X | X | | | X | X | | |
| 11/8 | **DPE 25** | X | X | X | X | X | X | X | X | | | | X | | |
| 11/11 | **DPE 28** | X | X | X | X | X | X | X | X | | | X | X | | |
| 11/15 | **DPE 32** | X | X | X | X | X | X | X | X | | | | X | | |
| 11/18 | **DPE 35** | X | X | X | X | X | X | X | X | | | X | X | | |
| 11/22 | **DPE 39** | X | X | X | X | X | X | X | X | | | | X | | |
| 11/29 | **DPE 46** | X | X | X | X | X | X | X | X | | | | X | | |

| | | | | | | | | | | | | | |
|---|---|---|---|---|---|---|---|---|---|---|---|---|---|
| 12/2 | **DPE 49** | X | X | X | X | X | X | X | X | | | | X |
| 12/6 | **DPE 53** | X | X | X | X | X | X | X | X | | | | X |
| 12/9 | **DPE 56** | X | X | X | X | X | X | | X | X | | | X |
| 12/13 | **DPE 60** | NA | NA | NA | X | X | X | X | X | | | | B, D, F |
| 12/16 | **DPE 63** | NA | NA | NA | X | X | X | X | X | | | | B, D, F |
| 12/20 | **DPE 67** | NA | NA | NA | X | X | NA | X | X | | | | B, D |

**Table S3. Model rankings.** Models were ranked according to Akaike's Information Criterion adjusted for small sample sizes ($AIC_C$). Top models were considered those within $\Delta AIC_C < 2$ of the top ranking model (bolded). Akaike's weight per each model is indicated as $w_i$.

| *Response variable* | *Model* | *df* | *log likelihood* | *AICc* | $\Delta AIC_C$ | $w_i$ |
|---|---|---|---|---|---|---|
| Prevalence of infection | **flock type * mean index infection duration** | **6** | **-258.63** | **529.33** | **0.00** | **0.52** |
| | **mean index infection duration** | **4** | **-261.40** | **530.82** | **1.50** | **0.24** |
| | flock type + mean index infection duration | 5 | -261.37 | 532.78 | 3.45 | 0.09 |
| | flock type | 4 | -262.46 | 532.96 | 3.64 | 0.08 |
| | intercept only | 3 | -263.79 | 533.59 | 4.27 | 0.06 |
| Prevalence of pathology | **intercept only** | **3** | **-314.85** | **635.72** | **0.00** | **0.50** |
| | **mean index infection duration** | **4** | **-314.80** | **637.63** | **1.91** | **0.19** |
| | **flock type** | **4** | **-314.84** | **637.70** | **1.98** | **0.19** |
| | flock type + mean index infection duration | 5 | -314.79 | 639.62 | 3.90 | 0.07 |
| | flock type * mean index infection duration | 6 | -314.28 | 640.62 | 4.90 | 0.04 |
| Infection likelihood | **sex*flock type** | **5** | **-32.78** | **76.33** | **0.00** | **0.46** |
| | **intercept only** | **2** | **-36.57** | **77.29** | **0.98** | **0.29** |
| | **sex** | **3** | **-35.64** | **77.59** | **1.26** | **0.25** |

**Table S4.** Model fits for stochastic SIR using the single naïve flock with ongoing transmission from index birds (Flock e; see Figure 2 and Table 1), with data for all three naïve flocks (also presented in Table 3) for comparison. Better fitting models have a lower median and minimum discrepancy, measured as the sums of squares difference (residual sums of squares; RSS) between model realizations and the empirical flock trajectories of infected birds over time. Note that RSS values for these single-flock model fits are not directly comparable with values for model fits using all three flocks since different datasets are used. Parameters from dose response data were from a previously published study (Hawley et al. 2024) and experimental epidemic data were collected in the current study. Note that a coefficient of variation (CV) for susceptibility is not reported for homogenous models because these models do not allow for any variation in susceptibility. Sixty-six percent confidence intervals are reported for means and CVs. [A] Reported values are the value of mean susceptibility (and CV) from fitted dose response data parameters, which is not identical to the median of the bootstrapped confidence intervals. [B] Parameter for the homogenous dose response data model restricted between 0 and 1, therefore not directly comparable to the mean susceptibility of the heterogeneous model. [C] Dose response data confidence intervals are computed via 1000 bootstrapped samples with replacement from data, therefore not directly comparable to the credible intervals computed in this study.

| Treatment | Model | Number of parameters fitted (n) | Dataset for susceptibility parameter estimates | Mean susceptibility: median estimate (66 % credible intervals) | CV susceptibility median estimate (66 % credible intervals) | Median Discrepancy (RSS) between model and flock data | Minimum Discrepancy (RSS) between model and flock data |
|---|---|---|---|---|---|---|---|
| Naïve | Homogeneous susceptibility | 1 | Dose response data (Hawley et al. 2024) | 0.838[A,B] (0.708, 1.000)[C] | -- | 291.5 | 3 |
| | | | Data from flock e only (this study) | 0.096 (0.043,0.18) | -- | 33 | 4 |
| | | | Data from all 3 naïve flocks (this study) | 0.062 (0.033,0.10) | -- | 33 | 4 |
| | Heterogeneous susceptibility | 2 | Dose response data (Hawley et al. 2024) | 1.18[A] (1.039, 1.530)[C] | 0.899[A] (0.805, 0.961)[C] | 60 | 4 |
| | | | Data from flock e only (this study) | 0.092 (0.035, 0.193) | 1.59 (1.09, 2.29) | 32 | 5 |
| | | | Data from all 3 naïve flocks (this study) | 0.06 (0.022, 0.088) | 1.79 (1.28, 2.44) | 32 | 4 |

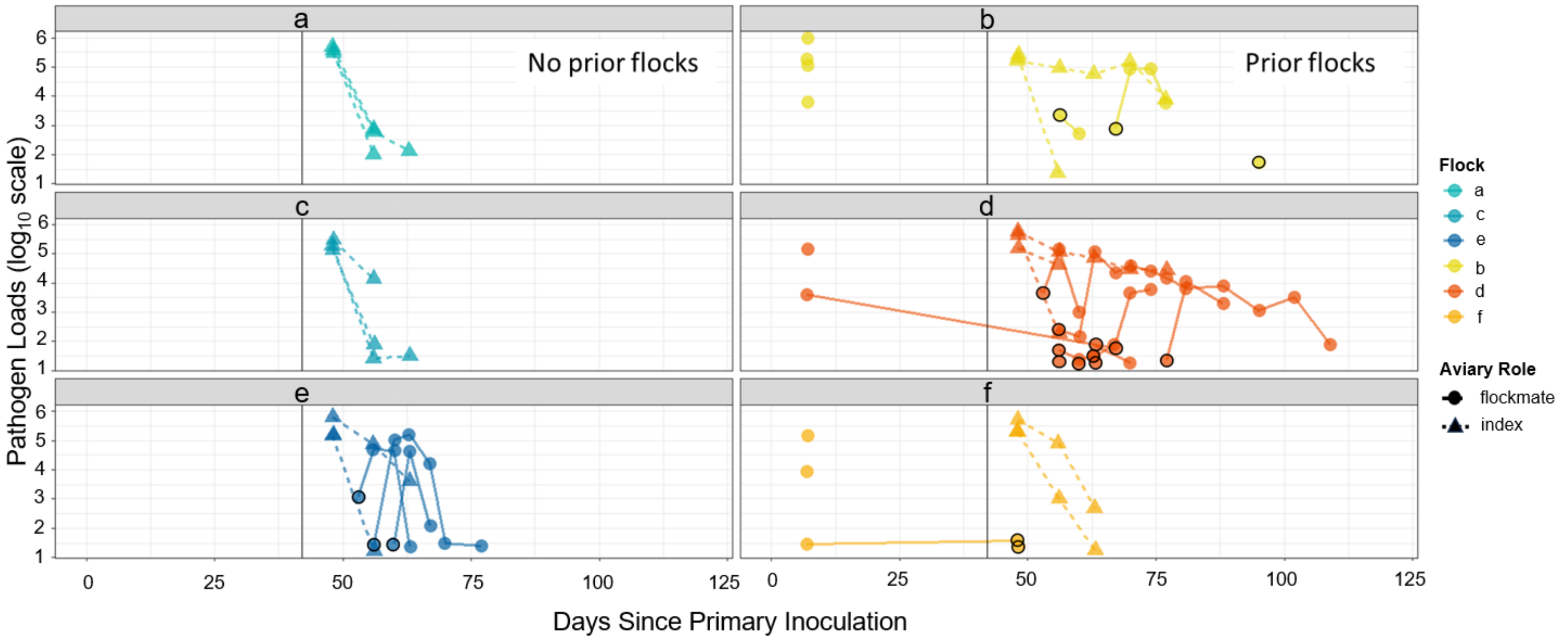

**Figure S1**. Each line segment or point represents a single bird (index [triangles/dashed lines] and flockmates [circles/solid lines]) at timepoints when it was infected. Only birds that showed pathogen load at some point during the study are included here. Left panels show naïve flocks (a, c, e) and the right panels show prior exposure flocks (b, d, f). Points outlined in black indicate the first epidemic infection timepoint per flockmate. The vertical line in each facet indicates the start of the epidemics. Birds that were infected both during their prior exposure (left of the vertical line) and in the epidemics were confirmed to be recovered and uninfected before the epidemics began but are shown with a connecting line here for visualization. Most likely, index bird infection duration was even more extreme than what we report or show here: although index bird eyeswabs were not systematically collected after epidemic day 35, opportunistic samples were collected on days 49 and 63 post-epidemic when two index birds were still actively infected. These samples had pathogen loads of $7.7 \times 10^4$ and $4.3 \times 10^3$, respectively.

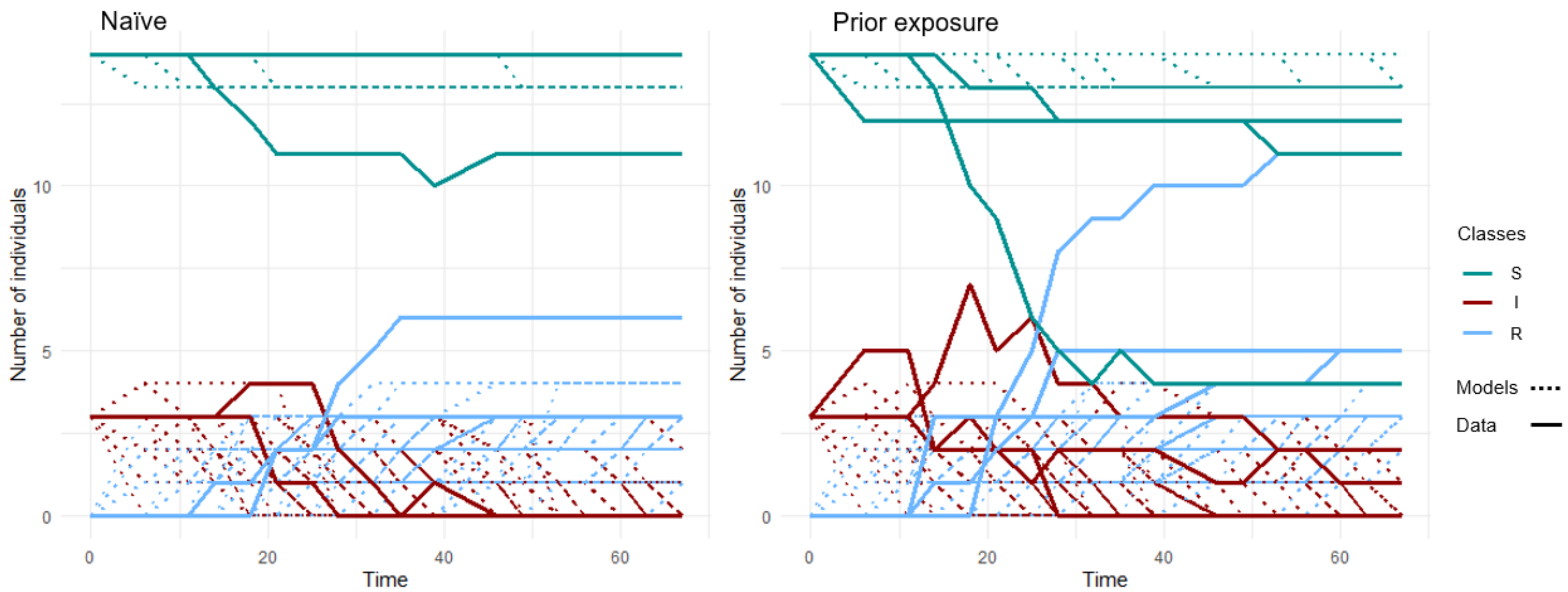

**Figure S2.** Stochastic model realizations (dotted lines; n = 100 realizations) and empirical data (solid lines) of an SIR model with heterogeneity in susceptibility, using best-fitting parameters for experimental flocks that were pathogen-naïve (left panel) or had prior exposure (right). This figure presents the same results as Figure 5 in the main text but includes the S and R classes in addition to the I class alone. Heterogeneity was described as a gamma distribution (naïve flocks: $k = 0.312$, $\theta = 0.129$; prior exposure flocks: $k = 0.08$, $\theta = 0.783$). Transmission ($\beta = 0.00275$) and recovery rates ($\gamma = 0.03$) were from prior literature (Hawley et al. 2024; Williams et al. 2014). Mortality $\alpha$ was set to zero because infection-induced mortality did not occur in captivity.
